# Single Cell Transcriptomics of Refractory Epilepsy patients in Colombia

**DOI:** 10.64898/2026.03.09.710691

**Authors:** Jorge Diaz-Riaño, Juan Pablo Carvajal-Dossman, Laura Guio, Daniel Mahecha, Paula Siaucho, Jennifer Guzman-Porras, Melissa Robles, Lina Bejarano, Paula Guzmán-Sastoque, Danilo Garcia-Orjuela, Andres Naranjo, Oscar Zorro, Silvia Maradei, Jorge Duitama

## Abstract

Maintaining electrical signaling homeostasis in the human neocortex relies on cell-type specific gene expression programs. However, when these programs are disrupted, the resulting imbalances can contribute to the pathogenesis of neurological disorders like epilepsy. Genetic factors are particularly implicated in a specific subtype of epilepsy known as refractory epilepsy or drug-resistant epilepsy (RE/DRE). This study shows the main results of the analysis of single cell transcriptomics for five pediatric RE patients in Colombia. A total of six samples obtained through surgical resection were analyzed by single-nuclei RNA sequencing (snRNA-seq). The genome of one patient was sequenced using high fidelity long-read sequencing. Functional enrichment of differentially expressed genes (DEGs) revealed glia-driven dysregulation of synaptic signaling, impaired glial–neuronal communication, and altered expression of genes related to neurotransmitter transport and calcium signaling. Activation of taste receptors in neurons was associated with neuroinflammatory processes. Structural variants were detected in genes associated with alterations of expression in specific cell types. This new data resource increases the diversity of information needed to develop new strategies for diagnosis of refractory epilepsy.

## INTRODUCTION

The human neocortex is a highly complex structure composed of tens to hundreds of distinct cell types, characterized by specific gene expression signatures (Jorstad et al., 2023). Gene expression regulates the production, differentiation, migration, and maturation of neuronal and glial cells. Disruptions in these genetic circuits can contribute to the pathogenesis of neurological disorders, including epilepsy, autism, Alzheimer’s disease, and Parkinson’s disease, among others (Velmeshev et al., 2023). Epilepsy is marked by recurrent episodes of brief yet intense bursts of electrical activity in the brain, known as seizures. These seizures can be focal (partial), affecting a specific brain region, or generalized, involving widespread neural networks and leading to significant behavioral changes (Milligan, 2021).

The International League Against Epilepsy (ILAE) defines epilepsy based on three diagnostic criteria: i) two or more unprovoked seizures occurring more than 24 hours apart, ii) one unprovoked seizure with a high risk of recurrence (greater than 60% according to additional studies) over the next 10 years, or iii) a diagnosis of an epilepsy syndrome even if seizures have not occurred yet (Fisher et al., 2014). Epilepsy is a prevalent condition affecting individuals of various age groups, particularly common before the age of 1 year and after the age of 50 (Kwan & Brodie, 2010). In Colombia, epilepsy affects 11.3 individuals per 1,000, with an incidence rate of 81.7 per 100,000 person-years, leading to an estimated loss of 5.25 disability-adjusted life years per thousand person-years (Orozco-Hernández et al., 2023).

Most epilepsy cases can be controlled with antiepileptic drugs (AEDs). However, medication alone is insufficient to prevent seizures in nearly a third of cases (Lerche, 2020). According to the International League Against Epilepsy (ILAE), epilepsy is diagnosed as drug-resistant (DRE) or refractory epilepsy (RE), when at least two appropriate AED regimens (as monotherapies or in combination), fail to achieve sustained seizure freedom (Kwan et al., 2010). In literature, ‘medically refractory’, or ‘pharmacoresistant epilepsy’ are used to describe the same condition. Several factors are associated with the risk of progression to DRE in patients with newly diagnosed epilepsy, including epilepsy onset in infancy, intellectual disability, symptomatic epilepsy, and abnormal neurological examinations. For some patients with RE, surgical resection, which involves seizure focus removal or brain activity-regulating device implantation (Guery & Rheims, 2021) may be considered. However, depending on factors such as the type and location of the intervention, among others, surgical treatment can be successful in approximately 40% to 80% of cases (Hsieh et al., 2023).

Although epilepsy can be caused by structural brain abnormalities and metabolic disorders, nearly 30-40% of epilepsy cases are considered genetic (Rastin et al., 2023). In these genetic epilepsies, there is a reproducible association between a genetic variant and the epilepsy phenotype (Scheffer et al., 2017). For instance, in genetic generalized epilepsies, which account for 15-20% of all epilepsies, a missense mutation in GABRG2, encoding the γ-aminobutyric acid (GABA A) receptor γ2 subunit, has been associated with childhood absence epilepsy (Hernandez et al., 2021). Additionally, a pathogenic missense variant in another GABA A receptor gene, GABRA1, encoding the α1 subunit, has been identified in myoclonic epilepsy (Perucca et al., 2020). For focal epilepsies, which represent 60% of all epilepsy cases, recent research reported significant genetic factors, particularly involving ion-channel genes (Perucca, 2018). Some examples include the voltage-gated potassium genes KCNQ2 and KCNQ3, linked to self-limited familial neonatal epilepsy; the voltage-gated sodium-channel α2 subunit gene SCN2A, associated with self-limited familial neonatal-infantile epilepsy; and the LGI1 gene, which encodes a neuronal secreted protein and is connected to autosomal dominant epilepsy with auditory features, among others (Perucca et al., 2020). Furthermore, genes encoding the subunits of the GATOR1 complex (DEPDC5, NPRL2, and NPRL3), which act as negative regulators of the mTOR signaling pathway, have been identified as having a significant role in this category of epilepsy (Baldassari et al., 2019; Balestrini et al., 2016). The association between genetic factors and proposed mechanisms of AEDs resistance has been described, and mainly includes alterations to the structure and/or functionality of brain targets of AEDs (such as voltage-dependent ion channels, neurotransmitter receptors, and transporters or metabolic enzymes involved in the release, uptake, and metabolism of neurotransmitters) (Löscher et al., 2020). For instance, splice-site or missense variants of the SCN1A gene can produce a reduced response in sodium channels, which provokes the blocking of AEDs through phenotypic effects like protein truncation, splice-site or missense variants. Another example is the inadequate penetration of antiepileptic drugs (AEDs) due to increased expression of efflux transporters, such as P-glycoprotein (Pgp), which was specifically observed in tissues obtained from resection surgery in DRE patients (Tang et al., 2017). This overexpression could reduce the bioavailability of AEDs even before these drugs arrive at the Blood Brain Barrier (BBB).

Advances in epilepsy research have been made in the past decade, particularly in the molecular genetics of the disease. The understanding of *de novo* mutations and the expression of genes (including regulators as miRNAs) has provided new insights about epileptogenesis. Through the use of single-cell RNA sequencing (scRNA-Seq) technology, researchers have identified distinct patterns of GABA_A receptor subunit expression changes in neuronal populations (such as pyramidal neurons and interneurons) and glial cells in samples from temporal lobe in animal models (Jorstad et al., 2023; Lee et al., 2023). Furthermore, a pro-inflammatory microenvironment within the epileptic lesions with an extensive activation of microglia and the infiltration of other proinflammatory immune cells were described by Kumar and collaborators (Kumar et al., 2022) applying CITE-Seq, a variation of scRNA-Seq.

This study aims to explore the cellular and molecular landscape of refractory epilepsy in Colombian patients. Single-cell gene expression profiling was performed for six samples taken from five different patients of refractory epilepsy. Differentially expressed genes (DEGs) were identified between these samples and appropriate controls for both neuronal and glial cells. One of the patients was also subject to long read whole genome sequencing to identify structural variants that could be related to DEGs or to genes previously related to epilepsy. This experiment allowed us to assess the power of long read sequencing to improve genetic diagnosis for epilepsy patients.

## RESULTS

### Cell type profiling of patients with refractory epilepsy

Single-nuclei RNA sequencing data was obtained from six brain samples (termed P010F, P010P, P018, P020, P024, and P025) belonging to five different patients **(Supplementary Table S1)**. Samples were extracted from frontal (including fronto temporal, and frontocentral), parietal and temporal regions of cortex. The sequencing yielded between 267 million (P010F) and 1.43 billion (P020) reads per sample. After filtering cells with extreme gene counts, the number of cells ranged from 4,043 (P018) to 7,281 (P024). The mean reads per cell spanned from 50,093 (P010F) to 257,152 (P025), while median UMI counts per cell ranged from 6,534 (P025) to 9,434 (P024). Median genes per cell varied between 2,508 (P025) and 3,295 (P024) (Supplementary Table S2).

The differences in sequencing depth reflect the fact that the samples were processed in two independent sequencing batches. After processing the first batch (P010F and P010P), we observed relatively low sequencing saturation (58.6% and 61.4%, respectively). These saturation values indicated that additional sequencing depth could improve UMI capture. Based on this assessment, the second batch of samples (P018, P020, P024, P025) was sequenced at higher depth. This decision improved sequencing saturation, reaching 86.5%, 91.3%, 72.1%, and 89.9% for the respective samples. Although the median number of genes was not affected by this decision, the distribution around this median value improved, especially for the samples P020, P024 and P025 **(Supplementary Figure S1)**.

Each cell was classified based on a correlation analysis between the gene expression profiles (nTPM) of the Human Protein Atlas (HPA) reference and raw UMI counts for 19,217 common protein coding genes **(Supplementary dataset 1)**. Cell types could be assigned reliably for 29,053 cells (83.5%) following this approach **(Figure 1)**. The dataset was predominantly composed of Oligodendrocytes (27.5% of the total cells), followed by Excitatory Neurons (19.1%) and Microglial Cells (15.1%). The remaining cell types were Astrocytes (14.8%), followed by Inhibitory Neurons (12.5%), and Oligodendrocyte Precursor Cells (OPCs, 9.8%). The remaining 334 cells were removed because they were classified as other non-brain cell types.

**Figure 1.**
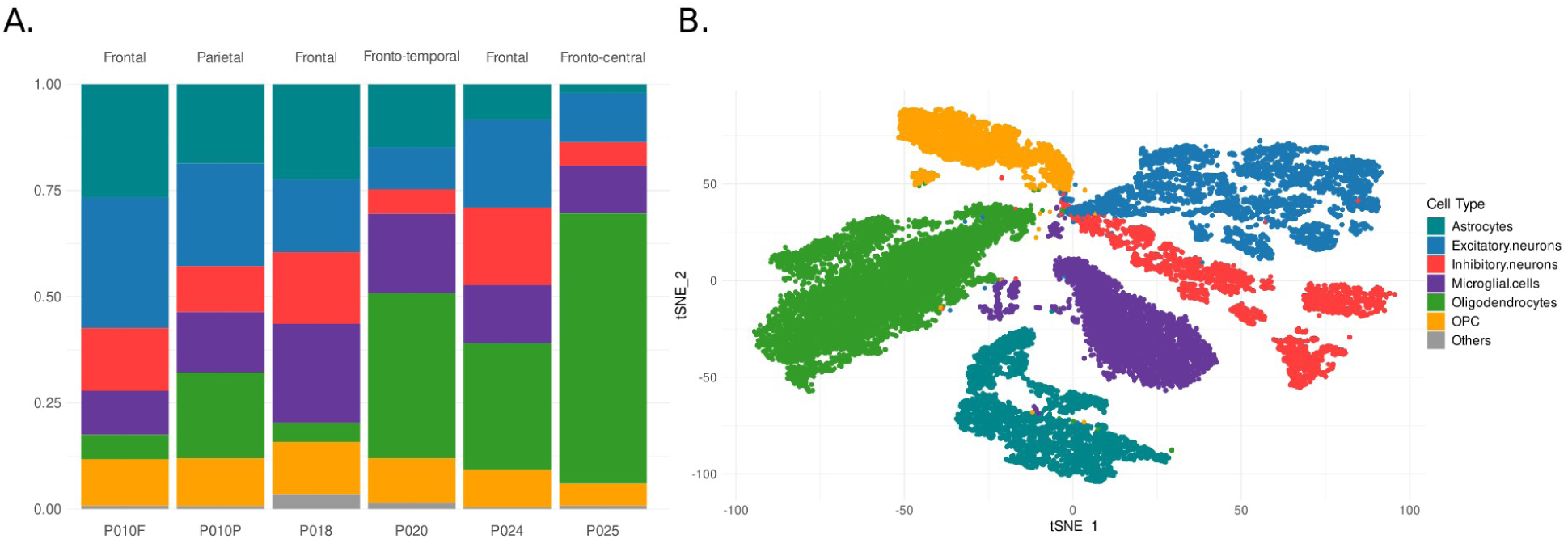
Cell type composition of refractory epilepsy dataset. **A.** stacked barplots per sample with origin regions for samples depicted on the bars. **B.** t-SNE of classified cell types in refractory epilepsy dataset.

A detailed analysis revealed differences in the composition of glial and neuronal cells across samples. Oligodendrocytes comprised ∼27.1% of the total cells per sample; the percentage of these cells within samples had a large variability, ranging from 4.5% (P018) to 63.51% (P025) accounting for a standard deviation of 22.3%. Astrocytes and microglial cells showed more consistent proportions across samples (∼15.5% and ∼15.3%, respectively), though astrocyte counts were notably lower in P024 and P025. Oligodendrocyte precursor cells (OPCs) were the most consistent glial category, representing ∼9.9% per sample, with a low standard deviation of 2.5%. Neuronal composition varied across samples, both in the ratio of inhibitory to excitatory neurons and in the overall neuron-to-glia ratio. Excitatory neurons accounted for ∼19.1% of total cells per sample (SD = 7.9%), while inhibitory neurons represented ∼11.96% (SD = 5.5%). The neuron-to-glia ratio ranged from 0.19 in P020 to 0.84 in P010F **(Figure 1)**.

### Differential expression in neuronal cells

To assess the impact of RE on different neuronal types, snRNA-seq data for 101,982 neuronal cells from healthy and affected donors analyzed in the study on Temporal Lobe Epilepsy (TLE) by Pfisterer et al. (2020) were compared with the expression patterns of the cells sequenced in this study. Cell type reassignment was cross-referenced with the original classification to establish a consistent set of excitatory and inhibitory neurons. These neuronal datasets were obtained from nine samples of healthy donors: Biopsy (C1), c_p1 (C2), c_p3 (C4), c_p4 (C5), GTS217_2 (C6), GTS219 (C7), GTS233 (C8), CTR215 (C9), and CTR240 (C10), all of which were designated as *controls*. An additional set of eight samples derived from patients affected by temporal lobe epilepsy (TLE), ep_p1 (E1), ep_p2 (E2), ep_p3 (E3), HB26 (E4), HB52 (E6), HB53 (E7), HB56 (E8), and HB65 (E9), comprised 22,464 excitatory and 16,976 inhibitory neurons and were used for cross-validation of cell-type assignments. These samples were obtained from an external published dataset and were designated as a reference TLE group (“Non-RE”). This label was used to distinguish them from the refractory epilepsy cohort generated in this study, but it does not imply a specific clinical classification.

Classification consistency (referring to the agreement between (i) the original cell-type labels provided in the external reference datasets and (ii) the cell-type labels assigned by our correlation-based annotation method) ranged from 89.4% (c_p4) to 97% (GTS233) in control samples and from 92.8% (HB53) to 97% (ep_p3) in Non RE samples, supporting the reliability of the reference-based cell type assignment method. The average composition of excitatory and inhibitory neurons was approximately 60% and 40%, respectively, across control, Non RE, and RE datasets **(Figure 2A)**. However, the control group exhibited the highest variance in the excitatory/inhibitory ratio, with values of 2.4, 0.6, 2.1, and 4.8 for Biopsy, c_p3, CTR240, and GTS233 samples, respectively. In the RE dataset, P018 and P024 showed the most distinct ratios. The RE dataset consisting of 9,108 neurons (5,509 excitatory and 3,599 inhibitory) was compared to 35,530 excitatory and 19,545 inhibitory reference neurons.

**Figure 2.**
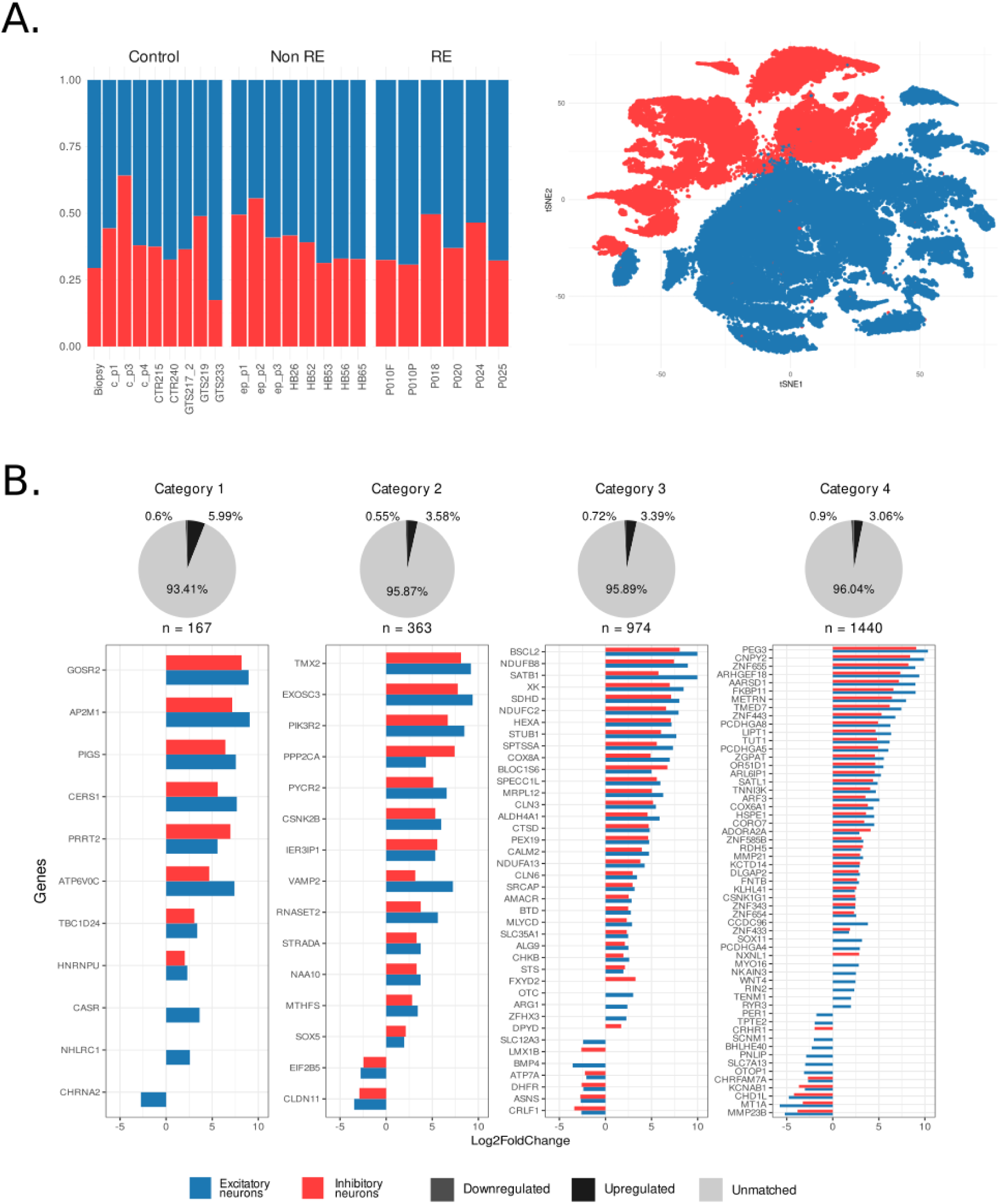
Neuronal dataset and expression. Cell-type composition across samples including TLE dataset. **A.** (left) Neuron-type composition for Control, Non refractory epilepsy, and refractory epilepsy datasets (right). **B.** Comparison of expression for epilepsy-associated genes in neurons. Pie charts depict the percentage of Upregulated/Downregulated genes matching against epilepsy-associated genes list (see methods section).

Comparison of pseudobulk gene expression values between control and RE neurons revealed a total list 1,304 differentially expressed genes (DEGs), 1,015 were upregulated, and 289 were downregulated in RE neurons **(Supplementary dataset 2)**. Excitatory neurons exhibited more DEGs (970 upregulated, 271 downregulated) than inhibitory neurons (776 upregulated, 163 downregulated) **(Supplementary Figure S2)**. A total of 731 upregulated and 145 downregulated genes were shared between both neuronal types, indicating that inhibitory neurons expressed fewer type-specific genes (45 upregulated, 18 downregulated) compared to excitatory neurons (239 upregulated, 126 downregulated).

Functional enrichment analysis using the Gene Ontology (GO) database identified a total of 23 significantly (adjusted p-value of a Fisher exact test ≤ 0.05) enriched categories **(Supplementary dataset 3)**. Among these, 17 categories (comprehending 12 biological processes and 5 molecular functions) were associated with upregulated genes, while 6 categories (4 biological processes and 2 molecular functions) were enriched among downregulated genes. Overall, a greater number of enriched categories were observed in excitatory neurons (13 for upregulated and 4 for downregulated genes list) compared to inhibitory neurons (4 upregulated and 2 downregulated).

GO enrichment based on upregulated genes predominantly highlighted terms for excitatory neurons related to cell–cell adhesion, particularly homophilic cell adhesion mediated by plasma membrane adhesion molecules, including members of the protocadherin (*Pcdh*) family. Ontologies associated with sensory perception (such as taste receptor *Tas* genes) were also enriched. The latter terms were also supported by enrichment of KEGG pathways. Among downregulated genes, metabolic pathways such as cellular glucuronidation and the uronic acid metabolic process were enriched in excitatory neurons (supported by KEGG enrichment), while terms related to oxidoreductase activity were recovered in inhibitory neurons **(Supplementary Figure S3).**

The DEGs identified in this analysis were compared to a list of 2,944 epilepsy-associated genes compiled by Zhang and colleagues (Zhang et al., 2024). This list categorizes genes into four groups: (1) 167 genes potentially associated with epilepsy as a primary or core symptom, (2) 363 genes linked to developmental disorders that include epilepsy, (3) 974 genes associated with other systemic abnormalities where epilepsy is a symptom, and (4) 1440 genes linked to epileptic phenotypes not included in the OMIM database but potentially related to epilepsy **(Supplementary dataset 4)**. A total of 123 DEGs overlapped with epilepsy-associated genes, distributed across four categories: 11 (∼9%) in category 1, 15 (∼12%) in category 2, 40 (∼33%) in category 3, and 57 (∼46%) in category 4 (see methods section- snRNA-seq and data analysis). All genes shared between excitatory and inhibitory neurons displayed consistent expression patterns. A total of 29 genes were cell type–specific, with 5 unique to excitatory neurons and 24 to inhibitory neurons. No genes exhibited opposite (i.e., inverse) differential expression profiles between the two neuronal types **(Figure 2B).**

### Differential expression in glial cells

The newly sequenced SCRNA data was also compared with public data from five donors (H18.30.001, H18.30.002, H19.30.001, H19.30.002, and H19.30.004) available at the Human Cell Atlas under the Supercluster of Non-Neuronal Cells (SCNNC) collection. These samples include a total of 108,638 cells. After quality filtering, cell type assignment was performed using the approach described in the methods section, and only cells assigned to a cell type consistent with that annotated in the public dataset were retained for analysis.

Classification consistency ranged from 80% for H18.30.002 to 99% for H19.30.004. The final SCNNC dataset consisted of 91,918 cells where oligodendrocytes accounted for approximately 49% (SD=14.2), while astrocytes comprised ∼27% (SD=8.1). Oligodendrocyte precursor cells (OPCs) and microglial cells were the least represented, making up ∼13.8% (SD = 4.8) and ∼9.9% (SD = 4.9) of the dataset, respectively **(Figure 3)**.

**Figure 3.**
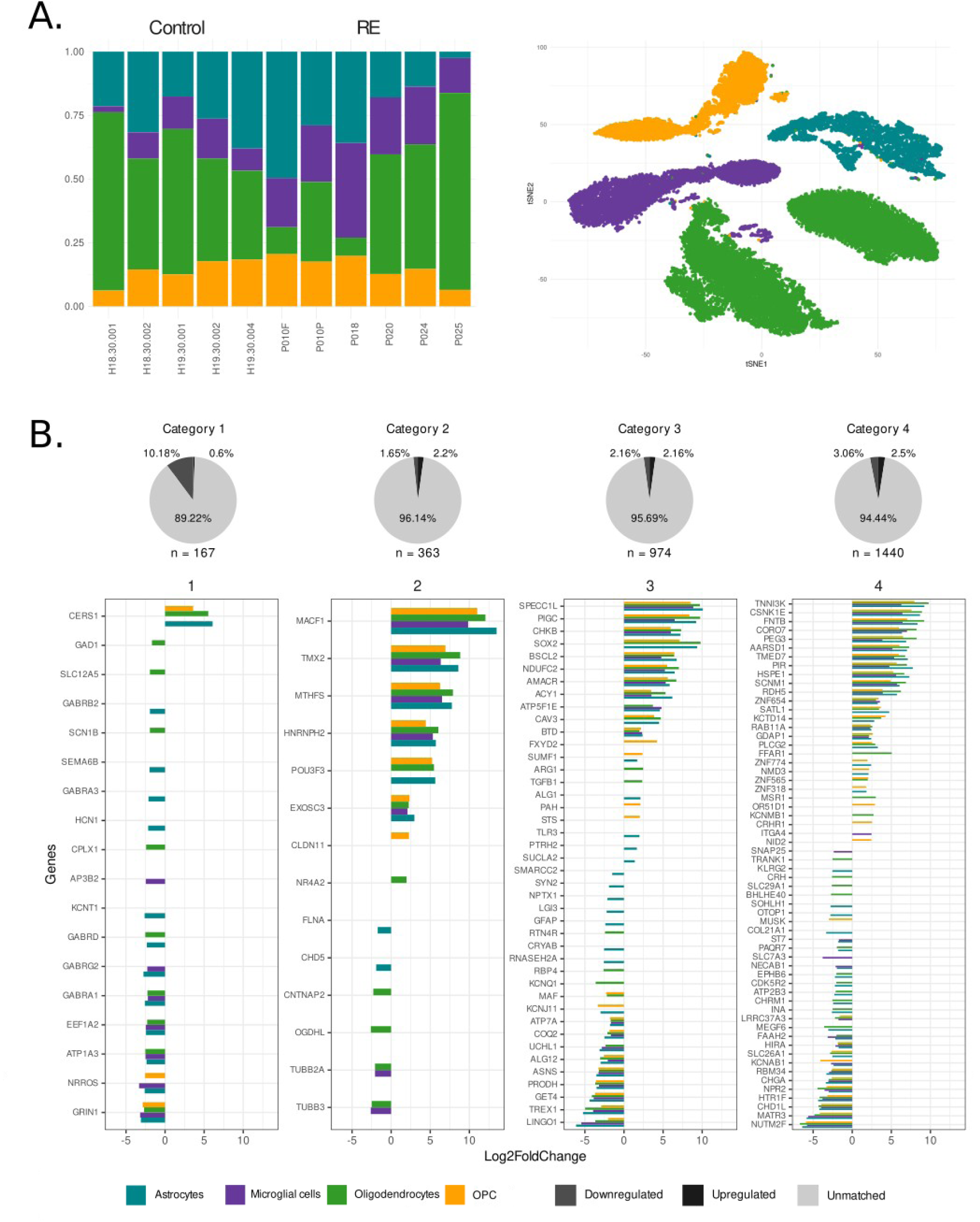
Glial composition and crossing with epilepsy-associated genes list. Cell-type composition across samples including the super cluster of non neuronal cells (SCNNC). **A.** (left) Glial cell-type composition for Control, and refractory epilepsy datasets. (right) t-SNE by glial cell-type. **B.** Comparison of expression for epilepsy-associated genes in glial cells. Pie charts depict the percentage of Upregulated/Downregulated genes matching against epilepsy-associated genes list. For visualization purposes, only the top 60 genes per category (ranked by absolute log2 fold change) were included in panel 4 (right).

A total of 1,160 genes were identified as differentially expressed (DEGs) across the cell types, where 602 were upregulated and 558 were downregulated **(Supplementary Figure S4, Supplementary dataset 2)**. Almost half of the DEGs (488) were shared across cell types (177 upregulated and 311 downregulated). The highest number of type-specific DEGs was found in Astrocytes (64 upregulated and 169 downregulated), and oligodendrocytes (53 upregulated and 77 downregulated). In microglial and opc cell types the numbers were similar (28, 32 upregulated, and 40, 25 downregulated respectively).

A total of 217 GO categories were enriched across all differentially expressed genes (DEGs) in glial cells, 206 associated with downregulated genes and 11 with upregulated genes **(Supplementary dataset 3)**. The majority of downregulated-enriched terms were found in astrocytes, including 65 biological processes, 46 cellular components, and 39 molecular functions. DEGs in oligodendrocytes enriched 14 biological processes, 19 cellular components, and 2 molecular functions. In microglial cells, 3 biological processes and 12 cellular components were enriched, while oligodendrocyte precursor cells (OPCs) accounted for 6 enriched cellular components. Most of the enriched terms were related to transporters, transmembrane ion flux (including potassium, calcium, and sodium), neurotransmitter transport, and modulation of synaptic signaling (particularly at glutamatergic and GABAergic synapses) with astrocytes and oligodendrocytes contributing to the majority of these functions **(Figure 4, Supplementary Figure S5).**

**Figure 4.**
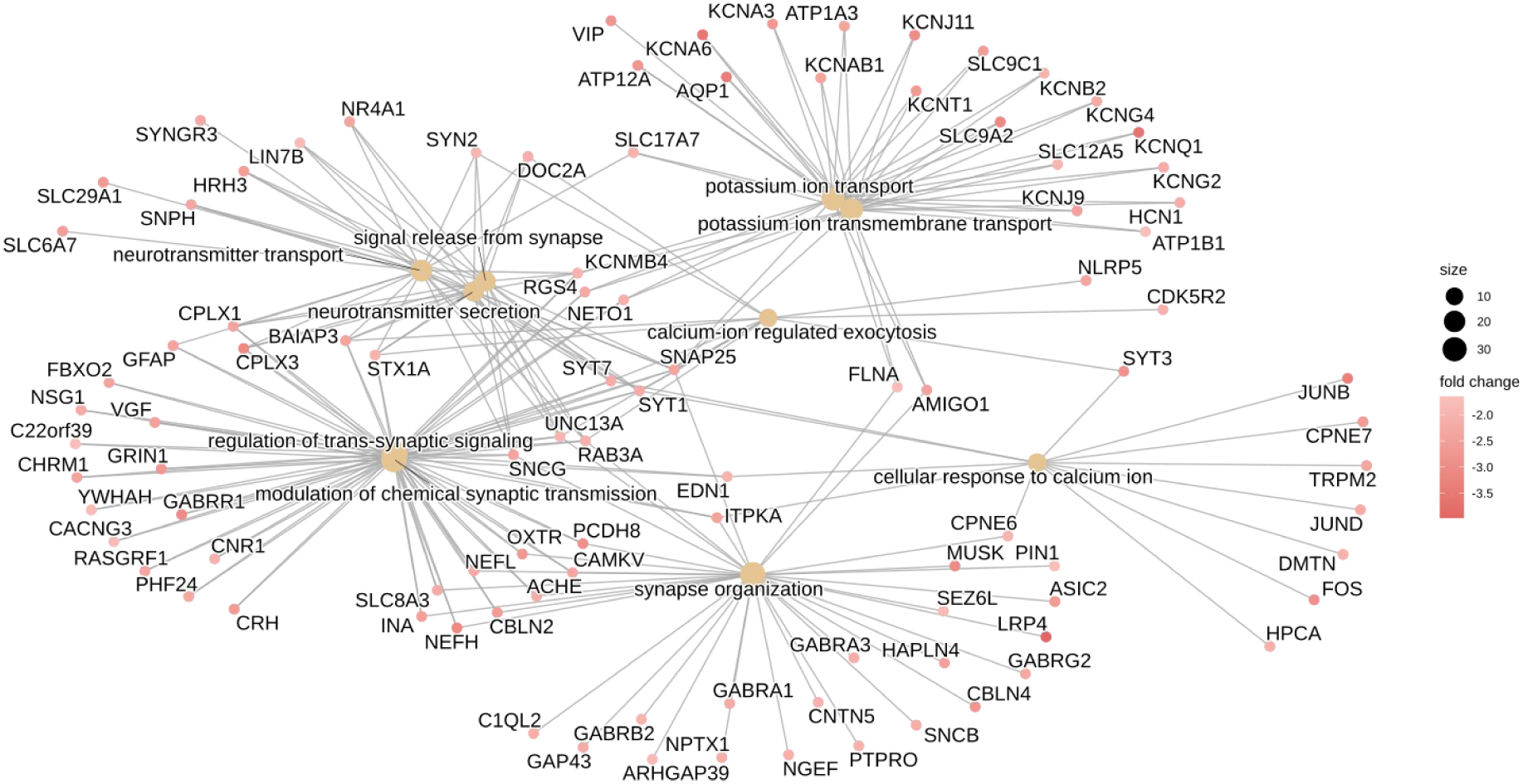
Functional Enrichment Glial Cells. Gene Ontology (GO) enriched terms with p.adjust ≤ 0.05 for downregulated genes in glial cells. The figure depicts the biological processes alongside related genes.

Regarding upregulated genes, the GO terms enriched in astrocytes included 2 cellular components and 4 molecular functions. Oligodendrocytes and OPCs exhibited 2 and 3 enriched molecular functions, respectively, while no enriched terms were identified for microglial cells. Among the enriched molecular functions were taste receptor activity, primarily involving *Tas* family genes; channel regulator activity, associated with *Slc* and *Kcn* gene families, both specifically enriched in OPCs; and the haptoglobin–hemoglobin complex, related to *Hb* family genes and enriched in astrocytes. **(Supplementary Figure S5).** Significant KEGG pathways related to human diseases (malaria-related signaling) and neuroactive ligand-receptor interaction were enriched in astrocytes and oligodendrocytes for downregulated genes. In oligodendrocytes, significant enrichment was observed for the taste transduction pathway showing a fold enrichment of 4.94 and involved several *Tas* family genes, reinforcing previous functional associations observed in these glial cells. MsigDB Molecular signatures (MSig) analysis highlighted TNF-alpha signaling via NF-κB and the Hallmark hedgehog signaling pathway in astrocytes.

A total of 154 DEGs from glial cells overlapped with previously reported epilepsy-associated genes. Of these, 18 genes (12%) matched category 1, 14 (9%) matched category 2, 42 (27%) matched category 3, and 80 (52%) matched category 4. Astrocytes and oligodendrocytes exhibited the highest number of matches, with 108 and 91 genes respectively, while OPCs and microglial cells showed fewer matches (75 and 66 genes, respectively). Similarly, the highest numbers of cell-type-specific genes (i.e., genes not expressed in other glial types) were found in astrocytes (32) and oligodendrocytes (20), followed by OPCs (13) and microglia (8). No inverse patterns of differential expression were found between cell types **(Figure 3B).**

### Genomic background of a refractory epilepsy patient

High fidelity long read whole genome sequencing was performed for one of the patients, aiming to obtain genomic information that could be related to refractory epilepsy. A read mapping approach followed by variant calling processes was performed to identify and genotype genomic variants relative to the reference genome. A total of 3,537,724 single nucleotide variants (SNVs) were identified. From these, 57.14% were genotyped as heterozygous. Consistent with the overall organization of the human genome and with conservation of exonic regions, more than 80% of the biallelic SNVs were present in intergenic and intronic regions. Only 24,800 SNVs were identified in protein coding exons, which accounts for an average of about one SNV per protein coding gene.

Following a two-tiered prioritization strategy for variant analysis based on the ACMG (American College of Medical Genetics and Genomics) variant classification guidelines (Richards et al., 2015), we identified a heterozygous missense variant in *SPEN* (NM_015001.3:c.3740G>T, p.Ser1247Ile, **Supplementary Figure S6A**). This SNV is absent from gnomAD, meeting the PM2 (Pathogenic Moderate 2: absent or extremely rare in population databases) criterion. Based on these criteria, the variant is classified as a variant of uncertain significance (VUS). Although it has not been associated with any disease to date, truncating variants in SPEN have been linked to a neurodevelopmental disorder characterized by developmental delay, intellectual disability, and other brain anomalies, including seizures in some individuals (Radio et al., 2021). Additionally, a heterozygous likely-pathogenic variant in *ACADS* (NM_000017.4:c.137G>A, p.Arg46Gln, **Supplementary Figure S6B**) was detected in a carrier state. This SNV results in an arginine-to-histidine substitution at codon 46 of the ACADS gene, which encodes the short-chain acyl-CoA dehydrogenase (SCAD) enzyme. This enzyme is crucial for mitochondrial fatty acid β-oxidation. The variant is rare in gnomAD, meeting the PM2 criterion , with multiple in silico predictive tools indicating a deleterious effect, meeting criterion PP3 (supporting evidence of pathogenicity: computational predictions consistent with a damaging impact on the gene or gene product). Additionally, the variant is a novel missense change at a residue where a different pathogenic missense change has been observed, fulfilling criterion PM5 (moderate evidence of pathogenicity: new missense change at an amino acid residue where another pathogenic missense change has been observed), and it has been reported as likely pathogenic in ClinVar, satisfying criterion PP5 (supporting evidence of pathogenicity: reputable source reports the variant as pathogenic). Based on these criteria, the variant is classified as likely pathogenic.

While the number and distribution of SNVs were consistent with previous works on human genetics (1000 Genomes Project Consortium, 2015), the use of long reads allowed us to identify a large and unbiased number of insertions and deletions relative to the reference genome. Combining algorithms to identify point mutations and structural variants, a total of 233,302 deletions and 237,411 insertions were identified. In contrast with SNVs, the percentage of these variants genotyped as heterozygous decreased to 23,63% for deletions and to 22,67% for insertions. Taking into account the potential impact of indels in protein translation, 97.10% of the indels were located in intergenic and intronic regions. The length distribution follows a power law with 89% of the indels having lengths between one and ten base pairs, and less than one thousand events spanning more than 1 Kbp **(Supplementary Figure S7A**). The longest variants included a homozygous deletion of 50,776 bp in the centromeric region of chromosome 16, and a heterozygous deletion of 21,744 bp in the centromeric region of chromosome 11.

Following the ACMG sequence variant interpretation guidelines, we identified a heterozygous variant of uncertain significance (VUS) of potential clinical interest: a 24 base pairs in-frame deletion in *GPRIN1* (NM_052899.3:c.693_716del, p.Glu233_Lys240del) **(Supplementary Figure S7B)**, resulting in the loss of eight amino acids (Glu233 to Lys240). It has a very low frequency in gnomAD (PM2), and it leads to the removal of multiple amino acids, potentially altering the function of the protein (PM4). Although this gene has not been associated with epilepsy or related disorders in OMIM, it encodes a protein that modulates G protein-mediated signal transduction at the cell membrane and within neuronal growth cones. GPRIN1 is involved in axonal and dendritic growth, cytoskeletal reorganization, and synapse formation, thereby regulating the architecture of the central nervous system (Nakata & Kozasa, 2005). This suggests that GPRIN1 can play a role in the pathogenesis of refractory epilepsy.

A total of 6,457 deletions affected 5,027 protein-coding genes **(Supplementary dataset 5)**. Of these, 567 are reported as epilepsy-associated genes: 42 as direct causes (category 1), 66 linked to developmental disorders that include epilepsy (category 2), 166 associated with conditions in which epilepsy is a symptom (category 3), and 293 currently under review as candidate genes (category 4). Similarly, 7,439 exonic insertions were related with 5,419 genes, including 625 epilepsy-associated genes: 32, 88, 195, and 310 for categories 1 to 4 respectively.

A total of 501 deletion and 508 insertion events overlapped with 616 DEGs identified by the analysis of scRNAseq data in at least one cell type (neuronal or glial). Only 90 of these DEGs were previously associated with epilepsy. In particular, a 3,169 bp heterozygous deletion was observed in the gene *RASA4* (chr7:102,588,788–102,591,957), resulting in a complete loss of the exon 17 **(Figure 5A)**. A prediction of tridimensional structure of the wild type and the alternative protein alleles (**Figure 5B**) shows that, although the overall structure remains largely conserved, the absence of the PH (Pleckstrin homology) domain induces a conformational change. PH domains direct proteins to specific cellular locations and facilitate interactions with binding partners, playing an important role in signal transduction pathways. Consequently, this alteration may impair normal protein function. When cross-referenced with cell-type–specific expression profiles, the gene was significantly upregulated in neurons. Although a direct link between this gene and epilepsy has not yet been established, *RASA4* encodes a member of the GAP1 family of GTPase-activating proteins that normally suppress the Ras/mitogen-activated protein kinase (MAPK) pathway in response to calcium signaling. Disruption of this regulatory function could alter downstream signaling dynamics in neurons, potentially shifting them toward a hyperexcitable state (Koh et al., 2018). The expression of *RASA4* was validated through RT-qPCR in a resection taken from an independent patient (Supplementary file 1).

**Figure 5.**
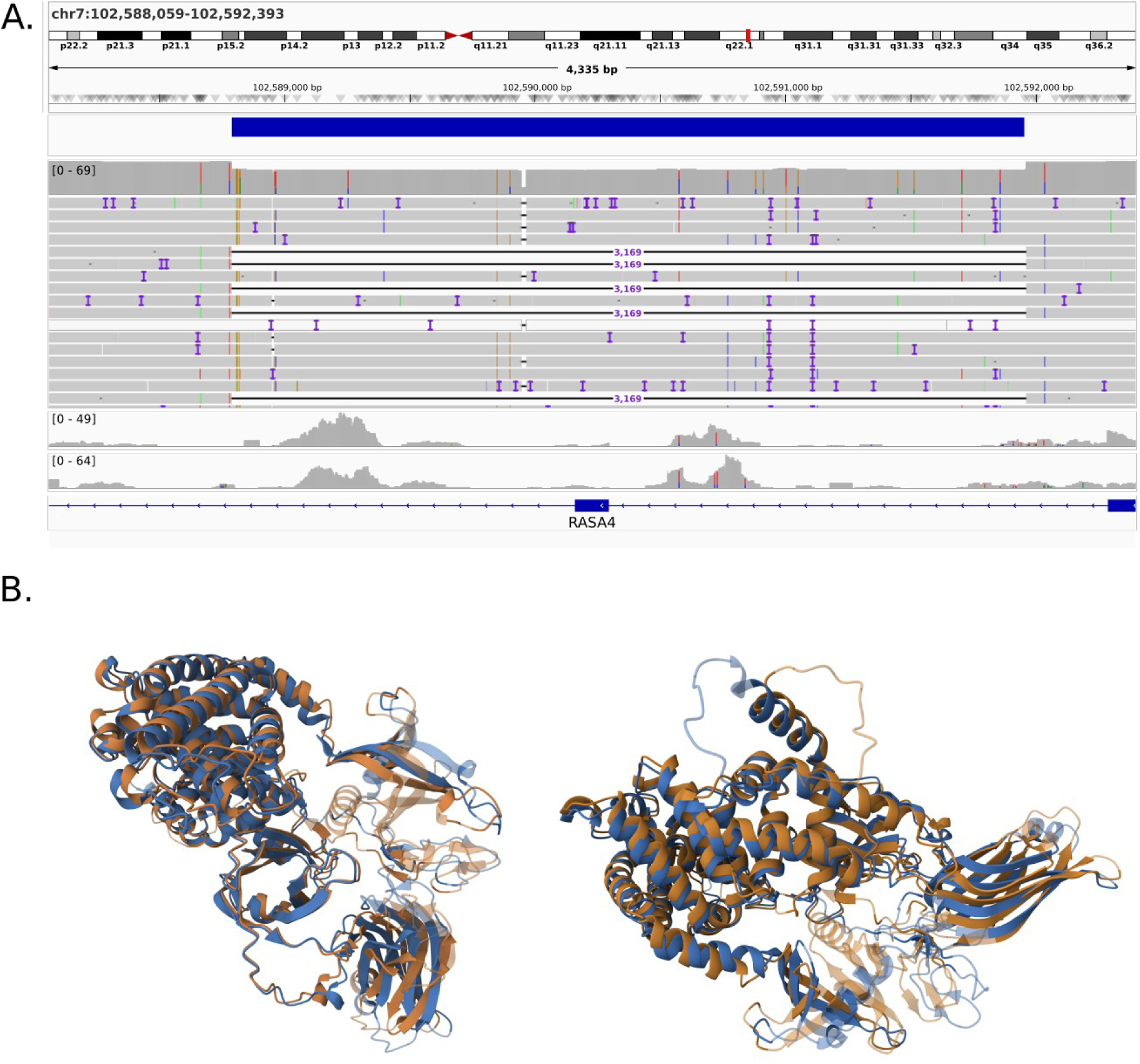
Structural variant associated with gene expression. A) Visualization in the Integrated Genome Viewer (IGV) showing a heterozygous deletion in the *RASA4* gene (potentially associated with the RAS pathway). The tracks at the bottom display the coverage of cDNA-mapped reads from excitatory and inhibitory neurons. B) The effect of a heterozygous structural deletion was modeled using AlphaFold. The figure shows two views of the structural alignment between the native protein (brown) and the altered protein with the deletion (blue). The faded region in the native structure corresponds to the segment absent in the altered form.

Another interesting variant is a 293 bp heterozygous insertion in the 3’ UTR of the gene *HCN1* (chr5:45,257,947–45,258,240), for which significant downregulation was observed in astrocytes. The expression of this gene was also validated through RT-qPCR (Supplementary file 1). In addition, 217 deletions and 464 insertions ranging from 1–100 bp in length affected genes previously related to epilepsy. For example, two variants exceeding 50 bp (a documented limit for short read sequencing) are a heterozygous deletion in the gene *KCNG2* (chr18:79,826,424 to 79,826,490) and a heterozygous deletion in the gene *KSR2* (chr12:117,465,752 to 117,465,808). In both cases, the genes were significantly downregulated in astrocytes.

## DISCUSSION

Epilepsy is a complex disease caused by a wide range of genetic and environmental factors. This work presents the main results of an exploratory study in which genomic and transcriptomic approaches were combined to obtain genetic and functional information on patients suffering from refractory epilepsy in Colombia. To the best of our knowledge, this is the first study providing expression information at single cell level from brain tissue in Latin America. Given the limited number of individuals that could be included in the study, the differential expression analysis was conducted taking as controls publicly available scRNA-seq datasets of “non disease” brain samples (autopsies of deceased individuals without known neurological disorders). We acknowledge that this experimental setting is far from ideal. However, even setting aside the economic constraints to generate snRNA-seq data in Colombia, it is still an open debate which samples should be used as controls to analyze expression data from samples with different neurological pathologies. Despite the limitations of the experimental design, the analysis revealed genes linked to expected functions such as synapses and signaling.

Although we initially expected a strong dysregulation affecting neuronal cells, the analysis revealed that glial cells had more clear alterations of synaptic transmission, compared to neurons. The observed downregulation of genes such as *BAIAP3*, *GAD2*, *GABRA1*, and *GABRG2* suggests a compromised ability to modulate synaptic function, sustain inhibitory signaling, and maintain neuroimmune balance. We hypothesize that glial cells actively participate in the promotion and maintenance of cortical hyperexcitability in refractory epilepsy (RE). Rather than being passive responders, glial cells are proposed to contribute to the convergence of multiple pathogenic mechanisms and to enable the emergence of alternative signaling routes that together sustain an epileptogenic environment, diminishing the effectiveness of conventional antiepileptic treatments. This hypothesis is supported by the recently introduced concept of "somatic microglial junctions", which refers to specialized contact points between microglial cells and neuronal soma (Cserép et al., 2020). These junctions are thought to enable microglia to monitor neuronal status via exocytosed signals, allowing for rapid immune and metabolic responses. Downregulation of *BAIAP3*, a gene implicated in vesicle docking and exocytosis, may disrupt this form of communication, hindering microglial capacity to detect neuronal distress or adjust to synaptic demands (Shen et al., 2023). These findings also align with growing evidence that glial cells not only provide structural and metabolic support to neurons but also engage in bidirectional communication that shapes the synaptic function and the dynamics of neuronal circuits (Shen et al., 2023). The contribution of glial cells to synaptic physiology is well-documented, particularly through processes such as neurotransmitter clearance and synaptic modulation. Astrocytes and oligodendrocytes participate in the process to remove glutamate and GABA from the synaptic cleft, regulating synaptic strength and preventing excitotoxicity (Bedner & Steinhäuser, 2023). In epileptic conditions, the downregulation of transporters involved in this function leads to prolonged excitatory signaling and reduced inhibitory tone, thereby increasing seizure susceptibility (Weston & Tzingounis, 2024). For instance, aberrant calcium signaling in astrocytes, and the consistent downregulation of a broad set of genes (including *BAIAP3*, *CPLX1*, *CRH*, *GABRA1*, *GABRR1*, *GRIN1*, *RAB3A*, *SYT7*, and *VGF*) can disrupt gliotransmitter release and metabolic support to neurons, further contributing to network instability and excitatory-inhibitory imbalance (Sumadewi et al., 2024), and potential impairments in axonal support and modulation of neurotransmission efficacy (Fields, 2015; Saab and Nave, 2017). The observed transcriptional changes may reflect a failure in microglial-neuronal communication, resulting in insufficient restraint of excitatory circuits. This could contribute to the persistence of seizure activity in refractory epilepsy by allowing unchecked excitability and impaired neuroimmune regulation. Future experiments can include immunohistochemistry or immunofluorescence to assess if glial alterations are related to refractory epilepsy.

One of the most interesting findings was the enrichment of upregulated taste receptors, particularly *TAS2R* genes (*TAS2R10, TAS2R13, TAS2R14, TAS2R30, TAS2R31, TAS2R4, TAS2R43, TAS2R46, TAS2R50,* and *TAS2R9*), in excitatory neurons. Although typically associated with chemosensory cells, *TAS2Rs* may have non-canonical roles in neurons (especially under pathological conditions such as epilepsy). Emerging evidence suggests that their abnormal activation can trigger neuroinflammatory responses, driven by increased production of proinflammatory mediators such as (cytokines and interleukins) and oxidative stress molecules (Welcome & Mastorakis, 2021). The expression of the gene *TAS2R14* was validated through RT-qPCR in a resection taken from an independent patient (Supplementary file 1). Although the metabolic pathways related to this neuroinflammatory response were not enriched in the analysis, genes such as *NLRP7* and *TNFSF12* showed a significant upregulation in excitatory neurons. The relationship between neuroinflammatory response and these genes have been reported recently (Wang et al., 2024). Future work can include secondary validation of these and other genes to assess if there is a causal or consequential relationship between these functions and refractory epilepsy.

We acknowledge that using public data as controls is prone to possible biases in the differential expression analysis caused by variables such as donor age, surgical methodology, and tissue origin. Age-related effects were considered when interpreting differential gene expression results. Steyn et al. (2024) identified several excitatory neuron genes that are expressed at higher levels in the pediatric cortex compared to adults. Three of these genes *(SOX11, TENM1*, and *MYO16*) were also upregulated in excitatory neurons in our dataset, indicating that a subset of the neuronal differential expression observed here may reflect developmental differences rather than disease-specific effects. In contrast, none of the non-neuronal age-associated genes reported by Steyn *et al*. were detected among the glial DEGs identified in this study. Additional age-associated genes reported in the literature, including *TNFRSF25* and *PIK3R3*, also showed differential expression in specific cell types in our dataset, further supporting the need to interpret neuronal findings in the context of developmental stage. Although formal age-matched controls were not available, the absence of age-associated signatures in glial DEGs and the convergence of glial pathways across patients suggest that the observed glial dysregulation is not solely attributable to developmental stage.

The region of the brain that was sampled during the surgery can be a main factor affecting the proportion of cells for each type. This variable completely depends on the epileptogenic focus for each patient and hence it can not be determined or controlled by the goals of the study. Taking this into account, the analysis pipeline considers possible batch effects confounding the analysis of the expression data. The relative composition of neuronal and glial cell types was similar across correspondent datasets. The main difference was observed in microglial cells, which were about 10% more abundant in the RE dataset for all the patients, compared to the control datasets. This could be explained by a more active immune response in brain regions affected by epilepsy and it is consistent with a neuroinflammatory cellular environment (Kumar et al., 2022; Johansen et al., 2023). However, the differential expression analysis of microglial cells did not reveal upregulation of genes such as cytokines or interleukins. Other possible explanations could be related to the natural genetic differences in human cortical cell composition (Johansen et al., 2023) or age-related aspects of cortical development (Velmeshev et al., 2023). Despite these differences, the unsupervised clustering of cells combining RE samples and controls grouped cells by cell type and not by study **(Supplementary Figure S8)**. This outcome was encouraging as a quality check for the sequencing, cell assignment and gene quantification processes. Another caveat of the experimental design proposed to identify DEGs can not distinguish between genes affected only by refractory epilepsy and genes affected by general epilepsy. This issue is particularly challenging because patients with non-refractory epilepsy are not subjected to surgery and hence it is ethically impossible to obtain brain tissue for these patients under the same conditions in which we obtained samples for patients with refractory epilepsy.

This study also includes a nearly complete genomic profiling for one of the patients, using a long read DNA sequencing protocol. The use of accurate long reads enables an unbiased discovery and genotyping of insertions and deletions in range lengths up to 20 Kbp. We acknowledge that genomic data for only one patient is clearly insufficient to identify a novel genetic element related to epilepsy. Rather than aiming at novel gene discovery, the long-read sequencing analysis was designed to evaluate the added diagnostic and interpretative value of structural variant detection in a clinical epilepsy context. Although the ACMG guides provide sufficient information to classify the SNVs found in the genotyped individual, they are insufficient to perform a systematic analysis of the thousands of large insertions and deletions that are usually discovered using long read technologies. For that reason we highlighted structural variants according to their potential impact in relation with their size and their intersection with previously known genes or with DEGs identified in this study.

The analysis clearly characterized the variant (NM_052899.3:c.693_716del, p.Glu233_Lys240del). Although there is currently no clinical evidence linking this specific variant to any particular disease, G protein-coupled receptors have been associated with seizure generation and the development of epilepsy (Yu et al., 2019). Other interesting variants include missense SNVs in the genes *SPEN* and *ACADS*. Moreover, a large heterozygous deletion affected one of the exons of the *RASA4* gene. Recent insights suggest that even subtle, somatic changes in genes within the RAS pathway might influence neuronal networks in ways that promote epileptogenic activity (Sran & Bedrosian, 2023). These observations support the idea that genetic alterations affecting components of the RAS/MAPK pathway, such as *RASA4*, could contribute to the imbalance of signaling processes underlying seizure susceptibility. In addition, the insertion affecting the *HCN1* gene, which showed reduced expression in astrocytes, is consistent with previous studies reporting that *HCN1* downregulation is associated with increased neuronal hyperexcitability in epilepsy (Bleakley & Reid, 2024). Up until now, the relationship between genetic variation and gene expression has been examined primarily in neurons. However, the observed differential expression in astrocytes could suggest the presence of additional regulatory mechanisms that may modulate *HCN1* expression and function beyond neuronal contexts, potentially contributing to altered network excitability and increased seizure susceptibility.

Overall, the results obtained in this study should be interpreted as an exploratory, hypothesis-generating analysis rather than as a definitive identification of drug-resistant epilepsy–specific molecular signatures. In particular, the genomic analysis is exploratory taking into account that we do not have information at the population level to determine the allele frequencies of the variant alleles in patients with refractory epilepsy and non refractory epilepsy. Even as a genetic diagnosis experiment it should be treated cautiously, because there is still a lag in guidelines to perform genetic diagnosis of structural variants. However, the results suggest that accurate long reads can quickly become a viable alternative to perform genetic diagnosis, taking into account the amount of information of structural variation that can be obtained from these data and the possible causative roles of structural variants for epilepsy and other complex diseases.

## METHODS

### Patient selection and sequencing

Blood and tissue samples were collected from a cohort of patients receiving treatment from the Epilepsy Surgery, Neurosurgery, Neurology, and/or Genetics departments at Fundación Hospital de la Misericordia (HOMI) in Bogotá. Patients diagnosed with refractory epilepsy, per ILAE criteria, who were selected for neurosurgery procedure by the pediatric epilepsy team at HOMI were included. Patients with autoimmune diseases, diabetes mellitus, and cancer were excluded. The project was approved by the research ethics committee at HOMI, as evidenced by the minute 49 − 21 of the meeting held on June 1st of 2021. The samples were obtained as part of the prescribed examination procedure for the patient and following all applicable guidelines and regulations. Legal representatives of the patient signed an informed consent for research purposes and data were anonymized to avoid identification of the patients from the data. All ethical regulations relevant to human research participants were followed.

Peripheral blood for a 14 years old donor (P024), diagnosed with drug-resistant epilepsy was collected via standard venipuncture and stored at -80°C. High Molecular Weight (HMW) DNA extraction was performed using the Monarch HMW DNA Extraction Kit for Cells and Blood (NEB #T3050S), following the instructions of the manufacturer. 500 µL of pretreated blood from sample P024 was used, and DNA was eluted in 80 µL of the elution buffer. The sample was incubated overnight at room temperature for homogenization. For snRNA-seq, six samples corresponding to five different patients were removed and processed as independent samples. The defined epileptogenic regions for selected patients were focal and temporal, including frontotemporal and frontocentral regions in one case **(Table S1)**. Brain tissue samples from epileptogenic regions were collected intraoperatively, immediately placed on dry ice, and stored at −80 °C until further processing. Samples were processed at the Princess Margaret Genomic Centre. Nuclei were extracted from frozen tissue following the 10x Genomics Chromium 3’ v3.1 Nuc-Seq protocol (10x Genomics Chromium Single Cell 3’ Reagent Kits v3.1 User Guide). Approximately 7,000–7,500 nuclei per sample were captured using the 10x Chromium system, and libraries were generated using unique sample index barcodes according to the manufacturer’s instructions. Libraries were sequenced following standard 10x Genomics recommendations on an Illumina NovaSeq X platform, achieving an average sequencing depth of ∼150,000 reads per nucleus. FASTQ files were generated following standard quality control and processing pipelines.

### SnRNA-seq and data analysis

*Preprocessing:* Mapping and feature counting were performed using STARsolo v2.7.11a (Kaminow et al., 2021) against the GRCh38-2020-A human reference genome. Data was analyzed with Seurat v5.0.1 (Hao et al., 2024). Cells with mitochondrial content >5% and those with <100 or >10,000 expressed genes were excluded. Doublets were identified and removed through scDblFinder v3.15 package (Germain et al., 2021).

Cells were classified into neuronal (inhibitory/GABAergic or excitatory/glutamatergic), non-neuronal (astrocytes, microglia, oligodendrocyte precursor cells, and oligodendrocytes), or non–brain-related cell types. For each nucleus, Pearson correlations were computed between its raw UMI counts (19,217 coding genes) and the normalized expression profiles (nTPM) of the Human Protein Atlas (HPA) reference dataset (Karlson et al., 2021; Siletti et al., 2023). The cell type associated with the highest correlation value was assigned as the initial biological classification. This correlation-based approach was Implemented as described by Robles et al. (2024). The cell type assignment was also validated looking at the expression patterns for canonical marker genes associated with each cell type **(Supplementary figure S9)**.

Control cells were obtained reanalyzing two publicly available datasets including glial and neuronal cells respectively. In the case of glial cells, controls were recovered from the supercluster of non-neuronal cells (SCNNC) dataset published in the Human Cell Atlas (Jorstad et al., 2023). For neuronal cells, a public dataset was analyzed containing samples from both healthy cortex biopsies (Pfisterer et al., 2020) and brain resections from patients with Temporal Lobe Epilepsy (TLE). As a validation of the previously mentioned process to assign cell types, the original annotations of SCNNC and TLE cells were compared with a reclassification performed using the same correlation process against expression profiles from the Human Protein Atlas (HPA). Only consistently classified cells were retained for further analysis. Consistency was measured as the proportion of cells for which this predicted label matched the original annotation in the reference dataset. To enhance dataset homogeneity, 16,315 cells originating from the primary visual cortex in the SCNNC collection were excluded, as this region is located in the occipital lobe, whereas the RE cells sequenced in this study were taken from the frontal and temporal lobes. For the TLE dataset, only control cells were included for the expression analysis. T-SNE visualizations by patient and by group were performed to assess possible sample or group batch effects **(Supplementary figure S10).**

A pseudobulking strategy was used to identify differentially expressed genes (DEGs) across annotated cell types. The expression values from the same cell type were aggregated by sample (considered as replicates) summing gene-specific UMI counts across all cells assigned to each type. This procedure was repeated independently for astrocytes, microglial cells, oligodendrocyte precursor cells (OPC), and Oligodendrocytes. Only genes with non-missing data across all samples were retained. A principal component analysis was used to evaluate potential batch effects **(Supplementary Figures S8)**. For neuronal pseudobulk profiles, batch effects were corrected using the ComBat-Seq method (Zhang et al., 2020). Differential expression analysis was performed through the DeSeq2 algorithm (Love et al., 2014) using an experimental design including the group (batch) and cell type. A shrinkage estimation of log2 fold changes was applied to reduce overestimation for genes with low counts.

The Gene Set Enrichment Analysis was conducted using GO (Thomas et al., 2022), KEGG (Kanehisa et al., 2025), and MSigDB (Liberzon et al., 2015) to identify over- and under-represented gene sets that may reveal mechanisms underlying the epileptic phenotype. The complete analysis workflow, including quality control, cell type annotation, pseudobulk construction, and downstream analyses, is fully documented and available at the following GitHub repository: https://github.com/jidiaz/scRNASeq-RefractoryEpilepsy_Colombia.

### Genomic variation analysis

Library Preparation and PacBio HiFi Sequencing Library preparation for PacBio HiFi Sequencing was performed at the Arizona Genomics Institute using the HiFi SMRTbell Prep Kit v3.0 with size selection on the PippinHT system. Sequencing was performed on the PacBio Revio platform using Revio SMRT Cells and Binding Kit v3, achieving a minimum Quality Value (QV) of 20 and a targeted depth of 30X.

Reads were mapped to the GRCh38 human reference genome using Minimap2 (Li, 2018). SNVs were identified with DeepVariant (Poplin et. al., 2018), and SVs with NGSEP v4.3.1 (Gaitán & Duitama, 2024). Variants were annotated using Clinvar v20240917, dbnsfp v4.7a, 1kG, GnomAD v4.1, and RefGene, as implemented in Annovar (Wang et al., 2010) and Ensembl VEP (McLaren et al., 2016). To prioritize SNVs and indels potentially associated with the clinical presentation of the patient, a two-tiered variant filtering strategy based on the ACMG guidelines (Richards et al., 2015) was implemented. Initially, a genotype-driven approach was applied by selecting variants meeting the aforementioned quality criteria and fulfilling at least one of the following conditions: i) previous classification as Pathogenic or Likely Pathogenic in ClinVar, ii) CADD *Phred* score greater than 20, or iii) AlphaMissense prediction as Pathogenic or Likely pathogenic. These candidate variants were further assessed using the ACMG guidelines.

The ACMG guidelines provide a variant classification framework based on weighted sources of evidence for pathogenicity or benignity. Evidence criteria are grouped in codes based on strength, and include PM (Pathogenic Moderate) and PP (Pathogenic Supporting), among others. For example, PM2 refers to variants that are absent or extremely rare in large population databases, supporting a pathogenic role. PP3 indicates that multiple in silico prediction tools consistently support a deleterious effect on the gene function. PM5 is used when a novel missense change occurs at an amino acid residue where a different missense variant has already been established as pathogenic. PP5 denotes variants already reported as pathogenic by a reputable source. A full list of criteria and combination rules is provided in Richards et al., 2015.

Additionally, a phenotype-driven filtering approach was employed retaining variants with the aforementioned quality criteria and located in genes associated with “epilepsy” and “seizure” in the phenotype annotations. Variants were excluded if their allele frequency in gnomAD exceeded 3% or if they were already classified as Benign or Likely Benign in ClinVar. Missense, splice site, and indel variants were evaluated according to the ACMG criteria. For non-coding variants (i.e. intronic, 3’UTR, 5’UTR, upstream and synonymous) additional filters were applied, retaining variants with CADD phred scores greater than 15 (or not available) and gnomAD allele frequencies less than 1%. Finally, variants were screened against a Comprehensive Epilepsy Panel (1,057 genes). The prioritized variants were then assessed using the ACMG criteria (Richards, 2015). The pathogenicity of SNVs (filtered by QUAL > 40) and CNVs was assessed according to their respective ACMG/AMP guidelines (Richards et al., 2015; Riggs et al., 2020). CNV (Deletions and duplications) scoring was automated using *ClassifyCNV* (Gurbich et al., 2020), followed by an analysis of epilepsy-associated genes affected by the identified variations. For other structural variants (SVs), an in-house workflow was developed, incorporating annotation and evaluating potential phenotypic effects.

Counts of exonic variants, classified as homozygous or heterozygous and as synonymous or nonsynonymous, were obtained for each gene. These exonic variants were compared against DEG lists to identify relationships between genomic changes and potential effects on expression that could explain the phenotypic expression of RE. Structural predictions for interesting variations were generated using the AlphaFold Server for both the full-length protein and the resulting isoform expressing the variant. The predicted models were aligned using the TM-align algorithm on the RCSB PDB web server.

### Statistics and Reproducibility

The six samples collected from brain resections of five different patients were treated as replicates for differential expression analysis. The eight controls for neurons and five controls for glial cells were also treated as replicates. Separate differential expression analyses were performed for neurons and glial cells. Pseudobulk counts were obtained for each cell type adding individual cell counts according to the cell type assignment. The DeSeq2 R package was used to perform differential expression analysis (Love et al., 2014). This package implements a generalized linear model to determine for each gene the estimated change of expression that can be attributed to each condition, after considering confounding factors such as sequencing depth and overdispersion for low counts. DeSeq2 implements a Wald test to calculate the p-value of the null hypothesis of no change in gene expression. It also implements the Benjamini–Hochberg procedure to perform correction for multiple testing. Genes were classified as differentially expressed if they showed an absolute log2 fold change greater than 1 and a corrected p-value (padj) less than 0.01. The data displayed in the figures is available at the **Supplementary dataset 6**.

## DATA AVAILABILITY

Raw data, including FASTQ files for single-cell RNA sequencing and genome sequencing, as well as matrix count files in MTX format for single-cell data, have been deposited in the NCBI Gene Expression Omnibus (GEO) under accession number GSE302285 and in the Sequence Read Archive (SRA) under BioProject ID PRJNA1291721 and BioSample accession SAMN49975736. The experiments also reanalyzed publicly available data from various repositories. Specific databases, accession IDs for each dataset, and reference genomes are detailed in the Materials and Methods section.

## CODE AVAILABILITY

The Python script implementing the cell-type classification is available with the distribution of scRANGE (Robles et al., 2024, https://github.com/mvrobles/scRANGE). All bioinformatic analyses were performed using publicly available, open-source software. The scripts used are available in the GitHub repository at https://github.com/jidiaz/scRNASeq-RefractoryEpilepsy_Colombia.

## ACKNOWLEDGEMENTS

This work was supported by the research fund of the Colombian Ministry of Science, “Conv. para Proyectos CTEI en Salud en Medicina Personalizada e Investigación Traslacional. Mod 2,” through grant contract No. 897-2021, awarded to J.D. The authors also acknowledge the support of the IT Services Department and ExaCore—IT Core Facility of the Vice Presidency for Research & Creation at Universidad de los Andes, for providing HPC resources that have contributed to the research results reported in this paper.

## AUTHOR CONTRIBUTIONS

J.D., S.M and A.N conceived the study, procured funding and coordinated the project. O.Z., L.G., P.S., J.G.-P. and D.G.-O. collected the samples. P.S., J.D.-R, P.G.S. and S.M. performed lab work for DNA sequencing and for validation of gene expression through RT-qPCR. J.D.-R., J.P.C.-D., D.M., M.R., L.B. and J.D. performed bioinformatic analysis. J.D., S.M, L.G. and A.N. provided scientific guidance and interpretation of the results. J.D.-R, D.M., and J.D. drafted the manuscript. All authors reviewed and approved the latest version of the manuscript.

## REFERENCES

Baldassari, S., Picard, F., Verbeek, N. E., van Kempen, M., Brilstra, E. H., Lesca, G., et al. The landscape of epilepsy-related GATOR1 variants. Genetics in Medicine, 21(2), 398–408 (2019). 10.1038/s41436-018-0060-2

Balestrini, S., Milh, M., Castiglioni, C., Lüthy, K., Finelli, M. J., Verstreken, P., et al. TBC1D24 genotype–phenotype correlation: epilepsies and other neurologic features. Neurology, 87(1), 77–85 (2016). 10.1212/WNL.000000000000280

Bleakley, L. E., & Reid, C. A. HCN1 epilepsy: From genetics and mechanisms to precision therapies. Journal of Neurochemistry, 168(12), 3891–3910 (2024). 10.1111/jnc.15928

Bedner, P., & Steinhäuser, C. Role of impaired astrocyte gap junction coupling in epileptogenesis. Cells, 12(12), 1669 (2023). 10.3390/cells12121669

Boileau, C., Deforges, S., Peret, A., Scavarda, D., Bartolomei, F., Giles, A., et al. GluK2 Is a Target for Gene Therapy in Drug-Resistant Temporal Lobe Epilepsy. Annals of Neurology, 94(4), 745–761 (2023). 0.1002/ana.26723

Brooks-Kayal, A. R., Shumate, M. D., Jin, H., Rikhter, T. Y., & Coulter, D. A. Selective changes in single cell GABAA receptor subunit expression and function in temporal lobe epilepsy. Nature medicine, 4(10), 1166–1172 (1998). 10.1038/2661

Chen, S., Zhou, Y., Chen, Y., & Gu, J. fastp: An ultra-fast all-in-one FASTQ preprocessor. Bioinformatics, 34(17), i884–i890 (2018). 10.1093/bioinformatics/bty560

Chen, Y., & Wang, X. miRDB: An online database for prediction of functional microRNA targets. Nucleic Acids Research, 48(D1), D127–D131 (2020). 10.1093/nar/gkz757

Cheng, H., Jarvis, E.D., Fedrigo, O. et al. Haplotype-resolved assembly of diploid genomes without parental data. Nat Biotechnol 40, 1332–1335 (2022). 10.1038/s41587-022-01261-x

Cifuentes-Diaz, C., Canali, G., Garcia, M., Druart, M., Manett, T., Savariradjane, M., et al. (2023). Differential impacts of Cntnap2 heterozygosity and Cntnap2 null homozygosity on axon and myelinated fiber development in mouse. Frontiers in Neuroscience, 17, 1100121 (2020). 10.3389/fnins.2023.1100121

Cserép, C., Pósfai, B., Lénárt, N., Fekete, R., László, Z. I., Lele, Z.,et al. Microglia monitor and protect neuronal function through specialized somatic purinergic junctions. Science, 367(6477), 528-537 (2020). 10.1126/science.aax6752

Cutillo, G., Bonacchi, R., Cecchetti, G., Bellini, A., Vabanesi, M., Zambon, A., et al. Interstitial 6q deletion in a patient presenting with drug-resistant epilepsy and Prader-Willi like phenotype: An electroclinical description with literature review. Seizure: European Journal of Epilepsy, 109, 45–49 (2023). 10.1016/j.seizure.2023.05.01

Fisher, R. S., Acevedo, C., Arzimanoglou, A., Bogacz, A., Cross, J. H., Elger, C. E., et al. ILAE Official Report: A practical clinical definition of epilepsy. Epilepsia, 55(4), 475–482 (2014). 10.1111/epi.12550

Formenti, G., Abueg, L., Brajuka, A., Brajuka, N., Gallardo-Alba, C., Giani, A., et al. Gfastats: conversion, evaluation and manipulation of genome sequences using assembly graphs. Bioinformatics, 38(17), 4214–4216 (2022). 10.1093/bioinformatics/btac460

Friedländer, M. R., Mackowiak, S. D., Li, N., Chen, W., & Rajewsky, N. miRDeep2 accurately identifies known and hundreds of novel microRNA genes in seven animal clades. Nucleic Acids Research, 40(1), 37–52 (2012). 10.1093/nar/gkr688

Gaitán, N., & Duitama, J. A graph clustering algorithm for detection and genotyping of structural variants from long reads. GigaScience, 13, giad112 (2024). 10.1093/gigascience/giad112

Germain, P. L., Lun, A., Meixide, C. G., Macnair, W., & Robinson, M. D. Doublet identification in single-cell sequencing data using scDblFinder. F1000Research, 10 (2021). 10.12688/f1000research.73600.2

Hao, Y., Stuart, T., Kowalski, M. H., Choudhary, S., Hoffman, P., Hartman, A., et al. Dictionary learning for integrative, multimodal and scalable single-cell analysis. Nature biotechnology, 42(2), 293–304 (2024). 10.1038/s41587-023-01767-y

Hernandez, C. C., Tian, X., Hu, N., Shen, W., Catron, M. A., Yang, Y., et al. Dravet syndrome-associated mutations in GABRA1, GABRB2 and GABRG2 define the genetic landscape of defects of GABAA receptors. Brain communications, 3(2), fcab033 (2021). 10.1093/braincomms/fcac156

Hsieh, J. K., Pucci, F. G., Sundar, S. J., Kondylis, E., Sharma, A., Sheikh, S. R., et al. Beyond seizure freedom: Dissecting long-term seizure control after surgical resection for drug-resistant epilepsy. Epilepsia, 64(1), 103–113 (2023). 10.1111/epi.17445

Hodge, R. D., Bakken, T. E., Miller, J. A., Smith, K. A., Barkan, E. R., Graybuck, L. T., et al. Conserved cell types with divergent features in human versus mouse cortex. Nature, 573(7772), 61-68 (2019). 10.1038/s41586-019-1506-7

Huang, N., & Li, H. compleasm: a faster and more accurate reimplementation of BUSCO. Bioinformatics, 39(10), btad595 (2023). 10.1093/bioinformatics/btad595

Jin, S., Plikus, M. V., & Nie, Q. CellChat for systematic analysis of cell–cell communication from single-cell transcriptomics. Nature Protocols, 1–40 (2024). 10.1038/s41596-024-01045-4

Johansen, N., Somasundaram, S., Travaglini, K. J., Yanny, A. M., Shumyatcher, M., Casper, T., et al. Interindividual variation in human cortical cell type abundance and expression. Science, 382(6667), eadf2359 (2023). 10.1126/science.adf2359

Jorstad, N. L., Close, J., Johansen, N., Yanny, A. M., Barkan, E. R., Travaglini, K. J., et al. Transcriptomic cytoarchitecture reveals principles of human neocortex organization. Science, 382(6667), eadf6812 (2023). 10.1126/science.adf6812

Kaminow, B., Yunusov, D., & Dobin, A. STARsolo: accurate, fast and versatile mapping/quantification of single-cell and single-nucleus RNA-seq data. Biorxiv, 2021-05 (2021). 10.1101/2021.05.05.442755

Kanehisa, M., Furumichi, M., Sato, Y., Matsuura, Y., & Ishiguro-Watanabe, M. KEGG: biological systems database as a model of the real world. Nucleic Acids Research, 53(D1), D672–D677 (2025). 10.1093/nar/gkae909

Karlsson, M., Zhang, C., Méar, L., Zhong, W., Digre, A., Katona, B., et al. A single–cell type transcriptomics map of human tissues. Science advances, 7(31), eabh2169 (2021). 10.1126/sciadv.abh2169

Koh, H.Y., Kim, S.H., Jang, J., et al. *BRAF* somatic mutation contributes to intrinsic epileptogenicity in pediatric brain tumors. Nat Med 24, 1662–1668 (2018). 10.1038/s41591-018-0172-x

Kuleshov, M. V., Jones, M. R., Rouillard, A. D., Fernandez, N. F., Duan, Q., Wang, Z., et al. Enrichr: a comprehensive gene set enrichment analysis web server 2016 update. Nucleic acids research, 44(W1), W90–W97 (2016). 10.1093/nar/gkw377

Kumar, P., Lim, A., Hazirah, S. N., Chua, C. J. H., Ngoh, A., Poh, S. L., et al. Single-cell transcriptomics and surface epitope detection in human brain epileptic lesions identifies pro-inflammatory signaling. Nature Neuroscience, 25(7), 956–966 (2022). 10.1038/s41593-022-01095-5

Kwan, P., Arzimanoglou, A., Berg, A. T., Brodie, M. J., Allen Hauser, W., Mathern, G., et al. Definition of drug resistant epilepsy: consensus proposal by the ad hoc Task Force of the ILAE Commission on Therapeutic Strategies. Epilepsia 51(6), 1069–1077 (2010). 10.1111/j.1528-1167.2009.02397.x

Lamar, T., Vanoye, C. G., Calhoun, J., Wong, J. C., Dutton, S. B., Jorge, B. S., et al. SCN3A deficiency associated with increased seizure susceptibility. Neurobiology of disease, 102, 38–48 (2017). 10.1016/j.nbd.2017.02.006

Langmead, B., & Salzberg, S. L. Fast gapped-read alignment with Bowtie 2. Nature Methods, 9(4), 357–359 (2012). 10.1038/nmeth.1923

Langmead, B., Trapnell, C., Pop, M., & Salzberg, S. L. Ultrafast and memory-efficient alignment of short DNA sequences to the human genome. Genome Biology, 10(3), R25 (2009). 10.1186/gb-2009-10-3-r25

Lee, B. R., Dalley, R., Miller, J. A., Chartrand, T., Close, J., Mann, R., et al. Signature morphoelectric properties of diverse GABAergic interneurons in the human neocortex. Science, 382(6667), eadf6484 (2023). 10.1126/science.adf6484

Lerche, H. Drug-resistant epilepsy—time to target mechanisms. Nature Reviews Neurology, 16(11), 595–596 (2020). 10.1038/s41582-020-00419-y

Li, H. New strategies to improve minimap2 alignment accuracy. Bioinformatics, 37:4572–4574 (2021). doi:10.1093/bioinformatics/btab705

Liberzon A, Birger C, Thorvaldsdóttir H, Ghandi M, Mesirov JP, Tamayo P. The Molecular Signatures Database (MSigDB) hallmark gene set collection. Cell Syst. 1(6), 417–425. (2015). 10.1016/j.cels.2015.12.004

Löscher, W., Potschka, H., Sisodiya, S. M., & Vezzani, A. Drug resistance in epilepsy: clinical impact, potential mechanisms, and new innovative treatment options. Pharmacological reviews, 72(3), 606–638 (2020). 10.1124/pr.120.019539

Love MI, Huber W, Anders S. Moderated estimation of fold change and dispersion for RNA-seq data with DESeq2. Genome Biology 15, 550 (2014). doi:10.1186/s13059-014-0550-8.

Mahmud, M., Wade, C., Jawad, S., Hadi, Z., Otoul, C., Kaminski, R. M., et al. Translocator protein PET imaging in temporal lobe epilepsy: A reliable test-retest study using asymmetry index. Frontiers in Neuroimaging, 2, 1142463 (2023). 10.3389/fnimg.2023.1142463

McLaren, W., Gil, L., Hunt, S. E., Riat, H. S., Ritchie, G. R., Thormann, A., et al. The Ensembl Variant Effect Predictor. Genome biology, 17(1), 122 (2016). 10.1186/s13059-016-0974-4

Milligan, T. A. Epilepsy: A Clinical Overview. The American Journal of Medicine, 134(7), 840–847 (2021). 10.1016/j.amjmed.2021.01.038

Naito, E., Indo, Y., & Tanaka, K. Identification of two variant short chain acyl-coenzyme A dehydrogenase alleles, each containing a different point mutation in a patient with short chain acyl-coenzyme A dehydrogenase deficiency. The Journal of clinical investigation, 85(5), 1575–1582 (1990). 10.1172/JCI114607

Nakata, H., & Kozasa, T. Functional characterization of Galphao signaling through G protein-regulated inducer of neurite outgrowth 1. Molecular pharmacology, 67(3), 695–702 (2005). 10.1124/mol.104.003913

Orozco-Hernández, J. P., Marín-Medina, D. S., Valencia-Vásquez, A., Quintero-Moreno, J. F., Carmona-Villada, H., & Lizcano, A. Predictors of adverse effects to antiseizure drugs in adult patients with epilepsy from Colombia: A case–control study. Epilepsy & Behavior, 146, 109383 (2023). 10.1016/j.yebeh.2023.109383

Perucca, P. Genetics of focal epilepsies: what do we know and where are we heading?. Epilepsy currents, 18(6), 356–362 (2018). 10.5698/1535-7597.18.6.356

Pasquini, G., Arias, J. E. R., Schäfer, P., & Busskamp, V. Automated methods for cell type annotation on scRNA-seq data. Computational and Structural Biotechnology Journal, 19, 961–969 (2021). 10.1016/j.csbj.2021.01.015

Perucca, P., Bahlo, M., & Berkovic, S. F. The genetics of epilepsy. Annual review of genomics and human genetics, 21(1), 205–230 (2020). 10.1146/annurev-genom-120219-074937

Pfisterer, U., Petukhov, V., Demharter, S., Meichsner, J., Thompson, J. J., Batiuk, M. Y., et al. Identification of epilepsy-associated neuronal subtypes and gene expression underlying epileptogenesis. Nature communications, 11(1), 5038 (2020). 10.1038/s41467-020-18752-7 |

Poplin, R., Chang, PC., Alexander, D. et al. A universal SNP and small-indel variant caller using deep neural networks. Nat Biotechnol 36, 983–987 (2018). 10.1038/nbt.4235

Radio, F. C., Pang, K., Ciolfi, A., Levy, M. A., Hernández-García, A., et al. SPEN haploinsufficiency causes a neurodevelopmental disorder overlapping proximal 1p36 deletion syndrome with an episignature of X chromosomes in females. American journal of human genetics, 108(3), 502–516 (2021). 10.1016/j.ajhg.2021.01.015

Ranallo-Benavidez, T.R., Jaron, K.S. & Schatz, M.C. GenomeScope 2.0 and Smudgeplot for reference-free profiling of polyploid genomes. Nat Commun 11, 1432 (2020). 10.1038/s41467-020-14998-3

Rastin, C., Schenkel, L. C., & Sadikovic, B. Complexity in genetic epilepsies: a comprehensive review. International Journal of Molecular Sciences, 24(19), 14606 (2023). 10.3390/ijms241914606

Rhie A, Walenz BP, Koren S, Phillippy AM. Merqury: reference-free quality, completeness, and phasing assessment for genome assemblies. Genome Biology 21, 245 (2020). 10.1186/s13059-020-02134-9

Richards, S., Aziz, N., Bale, S., Bick, D., Das, S., Gastier-Foster, J., et al. Standards and guidelines for the interpretation of sequence variants: A joint consensus recommendation of the American College of Medical Genetics and Genomics and the Association for Molecular Pathology. Genetics in Medicine, 17(5), 405–424 (2015). 10.1038/gim.2015.30

Riggs, E. R., Andersen, E. F., Cherry, A. M., Kantarci, S., Kearney, H., Patel, A., et al. Technical standards for the interpretation and reporting of constitutional copy-number variants: A joint consensus recommendation of the American College of Medical Genetics and Genomics (ACMG) and the Clinical Genome Resource (ClinGen). Genetics in Medicine, 22(2), 245–257 (2020). 10.1038/s41436-019-0686-8

Robles, M., Díaz-Riaño, J., Forigua, C., Ojeda, S., Guio, L., Siaucho, P., et al. New algorithms for unsupervised cell clustering from scRNA-seq data. bioRxiv (2024). 10.1101/2024.11.22.624768

Rosch, R., Burrows, D. R., Jones, L. B., Peters, C. H., Ruben, P., & Samarut, É. Functional genomics of epilepsy and associated neurodevelopmental disorders using simple animal models: from genes, molecules to brain networks. Frontiers in cellular neuroscience, 13, 556 (2019). 10.3389/fncel.2019.00556

Scheffer, I. E., Berkovic, S., Capovilla, G., Connolly, M. B., French, J., Guilhoto, L., et al. ILAE classification of the epilepsies: Position paper of the ILAE Commission for Classification and Terminology. Epilepsia, 58(4), 512–521 (2017). 10.1111/epi.13709

Shen, W., Pristov, J. B., Nobili, P., & Nikolić, L. Can glial cells save neurons in epilepsy?. Neural Regeneration Research, 18(7), 1417–1422 (2023). 10.4103/1673-5374.360281

Siletti, K., Hodge, R., Mossi Albiach, A., Lee, K. W., Ding, S. L., Hu, L., Transcriptomic diversity of cell types across the adult human brain. Science, 382(6667), eadd7046 (2023). 10.1126/science.add7046

Sim, S. B., Corpuz, R. L., Simmonds, T. J., & Geib, S. M. HiFiAdapterFilt, a memory efficient read processing pipeline, prevents occurrence of adapter sequence in PacBio HiFi reads and their negative impacts on genome assembly. BMC genomics, 23(1), 157 (2022). 10.1186/s12864-022-08375-1

Sourbron, J., Jansen, K., Mei, D., Hammer, T. B., Møller, R. S., Gold, N. B., et al. SLC7A3: In Silico Prediction of a Potential New Cause of Childhood Epilepsy. Neuropediatrics, 53(01), 046–051 (2022). 10.1055/s-0041-1739133

Sran, S., & Bedrosian, T. A. RAS pathway: the new frontier of brain mosaicism in epilepsy. Neurobiology of Disease, 180, 106074 (2023). 10.1016/j.nbd.2023.106074

Steyn, C., Mishi, R., Fillmore, S., Verhoog, M. B., More, J., Rohlwink, U. K., et al. A temporal cortex cell atlas highlights gene expression dynamics during human brain maturation. Nature Genetics, 56(12), 2718–2730 (2024). 10.1038/s41588-024-01990-6

Sumadewi, K. T., de Liyis, B. G., Linawati, N. M., Widyadharma, I. P. E., & Astawa, I. N. M. Astrocyte dysregulation as an epileptogenic factor: a systematic review. The Egyptian Journal of Neurology, Psychiatry and Neurosurgery, 60(1), 69 (2024). 10.1186/s41983-024-00843-7

Tang, F., Hartz, A. M., & Bauer, B. Drug-resistant epilepsy: multiple hypotheses, few answers. Frontiers in neurology, 8, 301 (2017). 10.3389/fneur.2017.00301

Tasic, B., Yao, Z., Graybuck, L. T., Smith, K. A., Nguyen, T. N., Bertagnolli, D., et al. Shared and distinct transcriptomic cell types across neocortical areas. Nature, 563(7729), 72–78 (2018). 10.1038/s41586-018-0654-5

Tarailo-Graovac, M., & Chen, N. Using RepeatMasker to identify repetitive elements in genomic sequences. Current protocols in bioinformatics, 25(1), 4–10 (2009). 10.1002/0471250953.bi0410s25

Thomas, P. D., Ebert, D., Muruganujan, A., Mushayahama, T., Albou, L. P., & Mi, H. PANTHER: Making genome-scale phylogenetics accessible to all. Protein Science, 31(1), 8–22 (2022). 10.1002/pro.4218

Velmeshev, D., Perez, Y., Yan, Z., Valencia, J. E., Castaneda-Castellanos, D. R., Wang, L., et al. Single-cell analysis of prenatal and postnatal human cortical development. Science, 382(6667), eadf0834 (2023). 10.1126/science.adf0834

Vlachos, I. S., Zagganas, K., Paraskevopoulou, M. D., Georgakilas, G., Karagkouni, D., Vergoulis, T., et al. DIANA-miRPath v3.0: Deciphering microRNA function with experimental support. Nucleic Acids Research, 43(W1), W460–W466 (2015). 10.1093/nar/gkv403

Wang K, Li M, Hakonarson H. ANNOVAR: Functional annotation of genetic variants from next-generation sequencing data Nucleic Acids Research, 38, e164 (2010) 10.1093/nar/gkq603

Wang X, Xiong W, Li M, Wu L, Zhang Y, Zhu C, et al. Role of inflammatory cytokine in mediating the effect of plasma lipidome on epilepsy: a mediation Mendelian randomization study. Front. Neurol. 15:1388920 (2024). 10.3389/fneur.2024.1388920

Welcome, M. O., & Mastorakis, N. E. The taste of neuroinflammation: Molecular mechanisms linking taste sensing to neuroinflammatory responses. Pharmacological Research, 167, 105557 (2021). 10.1016/j.phrs.2021.105557

Weston, M. C., & Tzingounis, A. V. 45 potassium channels in genetic epilepsy: A functional perspective. In Jasper’s Basic Mechanisms of the Epilepsies (Oxford Academic) 921–952 (2024). 10.1093/med/9780197549469.003.0045

Yu, Y., Nguyen, D. T., & Jiang, J. G protein-coupled receptors in acquired epilepsy: Druggability and translatability. Progress in neurobiology, 183, 101682 (2019). 10.1016/j.pneurobio.2019.101682

Zhang, S., Li, X., Lin, J., Lin, Q., & Wong, K. C. Review of single-cell RNA-seq data clustering for cell-type identification and characterization. Rna, 29(5), 517–530 (2023). 10.1261/rna.078965.121

Zhang, M. W., Liang, X. Y., Wang, J., Gao, L. D., Liao, H. J., He, Y. H., et al. Epilepsy-associated genes: an update. Seizure: European Journal of Epilepsy, 116, 4–13 (2024). 10.1016/j.seizure.2023.09.021

Zhao, W., Johnston, K.G., Ren, H. et al. Inferring neuron-neuron communications from single-cell transcriptomics through NeuronChat. Nat Commun 14, 1128 (2023). 10.1038/s41467-023-36800-w

1000 Genomes Project Consortium. A global reference for human genetic variation. Nature, 526(7571), 68 (2015). 10.1038/nature15393

